# Label-free imaging of DNA interactions with 2D materials

**DOI:** 10.1101/2023.07.05.547763

**Authors:** Jenny Sülzle, Wayne Yang, Yuta Shimoda, Nathan Ronceray, Eveline Mayner, Suliana Manley, Aleksandra Radenovic

## Abstract

Two-dimensional (2D) materials offer potential as substrates for biosensing devices, as their properties can be engineered to tune interactions between the surface and biomolecules. Yet, not many methods can measure these interactions in a liquid environment without introducing labeling agents such as fluorophores. In this work, we harness interferometric scattering (iSCAT) microscopy, a label-free imaging technique, to investigate the interactions of single molecules of long dsDNA with 2D materials. The millisecond temporal resolution of iSCAT allows us to capture the transient interactions and to observe the dynamics of unlabeled DNA binding to a hexagonal boron nitride (hBN) surface in solution for extended periods (including a fraction of 10%, of trajectories lasting longer than 110 ms). Using a Focused Ion Beam (FIB) technique to engineer defects, we find that DNA binding affinity is enhanced at defects; when exposed to long lanes, DNA binds preferentially at lane edges. Overall, we demonstrate that iSCAT imaging is a useful tool to study how biomolecules interact with 2D materials, a key component in engineering future biosensors.

---

Two-dimensional (2D) materials have been developed as DNA sensing platforms owing to their chemical, electronic and mechanical properties which accommodate a variety of sensing modalities such as membrane-embedded nanopores^1–7^, tunneling junctions^8,9^ and field-effect transistors^10,11^. Such devices have succeeded in sensing individual DNA strands, both optically and electronically^12–16^. Different sensing modalities and functionalisations can be integrated to detect a whole host of biological analytes with a single device, which due to a high surface-to-volume ratio can be performed on limited sample volumes.

An important remaining challenge is to control surface interactions and to prevent fouling of the devices^17^. Toward this end, understanding DNA interactions with 2D materials is crucial to improve biosensor designs by reducing unwanted interactions and by promoting desired ones. A common approach is to label DNA fluorescently with intercalating dyes (such as YOYO-1) and study it with optical microscopy^18^. Alternatively, atomic force microscopy (AFM) can simultaneously detect molecular binding and map the surfaces^19,20^. Electron microscopy (EM) has also been used to image 2D materials in the presence of biomolecules at even higher spatial resolution^21–23^. However, each of these approaches suffers from limitations. Intercalating dyes change the length and stiffness of DNA^24,25^, and such dyes suffer from photobleaching and quenching near 2D material surfaces^26,27^. AFM offers imaging in liquid, but only for hydrophilic substrates such as mica. In the case of prototypical 2D materials, such as graphene, hexagonal boron nitride (hBN) or transition metal dichalcogenides (TMDCs), imaging is often limited to dried DNA on 2D material surfaces. EM is largely confined to dry specimens and thus lacks dynamic information, or requires special liquid holders or the use of gold nanoparticles as labels^28–32^. This may explain in part why simulation has led experiment in investigating interactions of DNA with more complex materials and device designs^33–36^.

In this work we harness interferometric scattering (iSCAT) microscopy^37,38^ to investigate single molecules of long double stranded (ds)DNA interacting with the surface of hBN. As an analyte in solution approaches the surface, interference between the light it scatters and the light reflected by the liquid-solid interface boosts the signal contrast, enabling single-molecule detection. Because iSCAT uses the molecules themselves to generate signal, it does not require labeling. Liberated from the limited photon budget of fluorescent dyes, the technique allows millisecond temporal resolution, and theoretically indefinite imaging times. So far, iSCAT has been used to study the mass distribution of short single-stranded and double-stranded DNA^39^, protein oligomers down to molecular weights of 10 kDa^40,41^, and the products of other biomolecular reactions^42^ as well as the dynamics of more complex cellular processes^43^. To study the effects of surface properties on DNA binding, we study its interactions with pristine hBN as well as engineered defects and nanostructures^44^.

We chose hBN as our substrate due to its optical properties: it is transparent because of its band gap of 6 eV^45^, and it uniquely among 2D materials^27,46–48^ lacks fluorescence quenching which allows integration with fluorescent sensing strategies. Additionally, we note the potential for future engineering^33^ due to its regular shaped crystal edges^49–51^ and single layer films which appear promising candidates for high-quality large-scale production^52–54^. hBN was exfoliated on coverslips and imaged with iSCAT (Figure 1a, SI Experimental Section). The hBN flakes show a static uniform scattering background when illuminated in water, with stronger scattering from their edges. To prevent edge scattering from overwhelming the weaker signal from dsDNA, we imaged the larger flakes, and ensured they filled the selected FOV (Figure S1). Furthermore, we use multiple frames to calculate a running averaged scattering background and to produce a differential contrast image.

**Figure 1.**
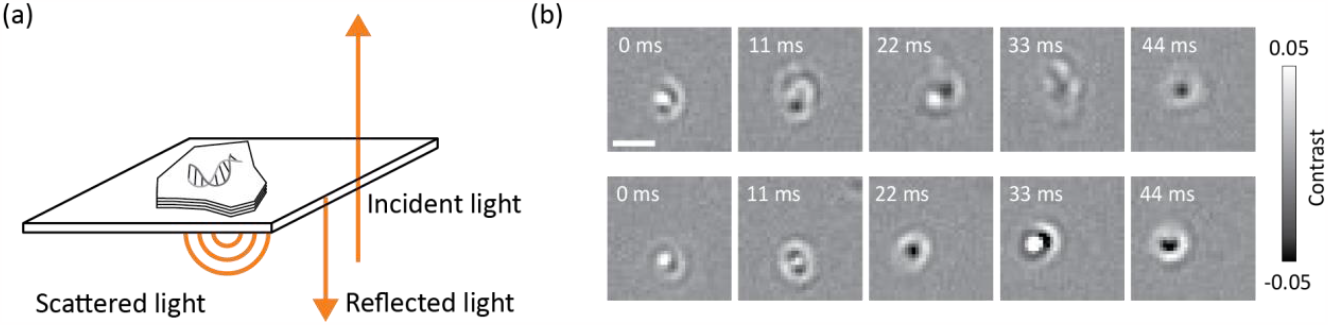
Interaction of 2D materials and DNA probed by iSCAT. (a) Schematic setup of the experiment. (b) Two examples of median-filtered time series of 20 kbp dsDNA particles diffusing on the hBN surface, illustrating changes in shape, size, and contrast of the same molecule from frame-to-frame. Scale bar is 1 µm.

Upon the addition of DNA, we observe the appearance of large scattering signals (blobs) with bright and dark regions in images following median-filtered background removal (Figure 1b). We measure similar patterns for a range of different DNA lengths: 10, 20, and 48 kbp (Figure S2). These sizes were chosen based on our observed lower detection limit of 10 kbp dsDNA. We expect that the observed blobs of DNA are of single molecules, consistent with observations of both labeled and unlabeled DNA by other single molecule sensing techniques employing similar concentrations (pM-nM) of DNA^55–57^.

Theoretical and experimental studies have demonstrated that iSCAT is highly sensitive to the axial position of scattering particles, and even nanometric changes can result in a contrast inversion in the temporally-filtered images, due to constructive and destructive interference^58,59^. These studies measured biomolecules much smaller than the diffraction limit, which led to compact, circularly shaped interferometric point spread functions (iPSF). In our case, the estimated radii of gyration for the measured DNA molecules are 238 nm (10 kbp dsDNA), 337 nm (20 kbp) and 521 nm (48 kbp). We speculate that the complex scattering signal is a superposition of multiple iPSFs from different parts of the same molecule, which spans several hundred nanometers axially. Similar studies with iSCAT show a linear correlation of the contrast signal to the molecular mass of the analyte for molecules much smaller than the diffraction limit^39,40,60^. Here, we chose the mean contrast signal of each localization to investigate its relationship to molecular weight, but we did not find a correlation for these large DNA molecules.

In time-lapse experiments, we qualitatively observe motion of the scattering blobs (Figure 1b, Figure S3 and Movie S1). The overall signal contrast is typically low in the frames when a particle approaches or leaves the surface, but it increases when particles are moving along the surface. We wished to quantify this motion, but we noticed that the same molecule could change contrast, shape, and size from frame to frame. We speculate that this is due to changing 3D conformations of the dsDNA on the hBN surface. Existing strategies for detection and localization such as thresholding and spot detection yield poor efficiency due to this complex appearance - this motivated us to develop a customized image analysis pipeline.

We developed an approach to segment and localize the particles for data with a temporal median-filtered background normalization (Figure 2a, Figure S4, S5, S6). We use the absolute value of the signal, to combine both constructive and destructive interference contributions to the signal. We optimize the threshold to avoid false positives from the noisy background, while still detecting faint signals. Each particle’s absolute contrast-weighted center of mass defines its location, which can be connected in time to reconstruct trajectories (SI Experimental Section). We manually validated the automatically-tracked results by checking each step in the pipeline for a randomly selected subset of particles. We define each detected localization as a DNA binding site, and the total time of reconstructed trajectories as dwell time.

**Figure 2.**
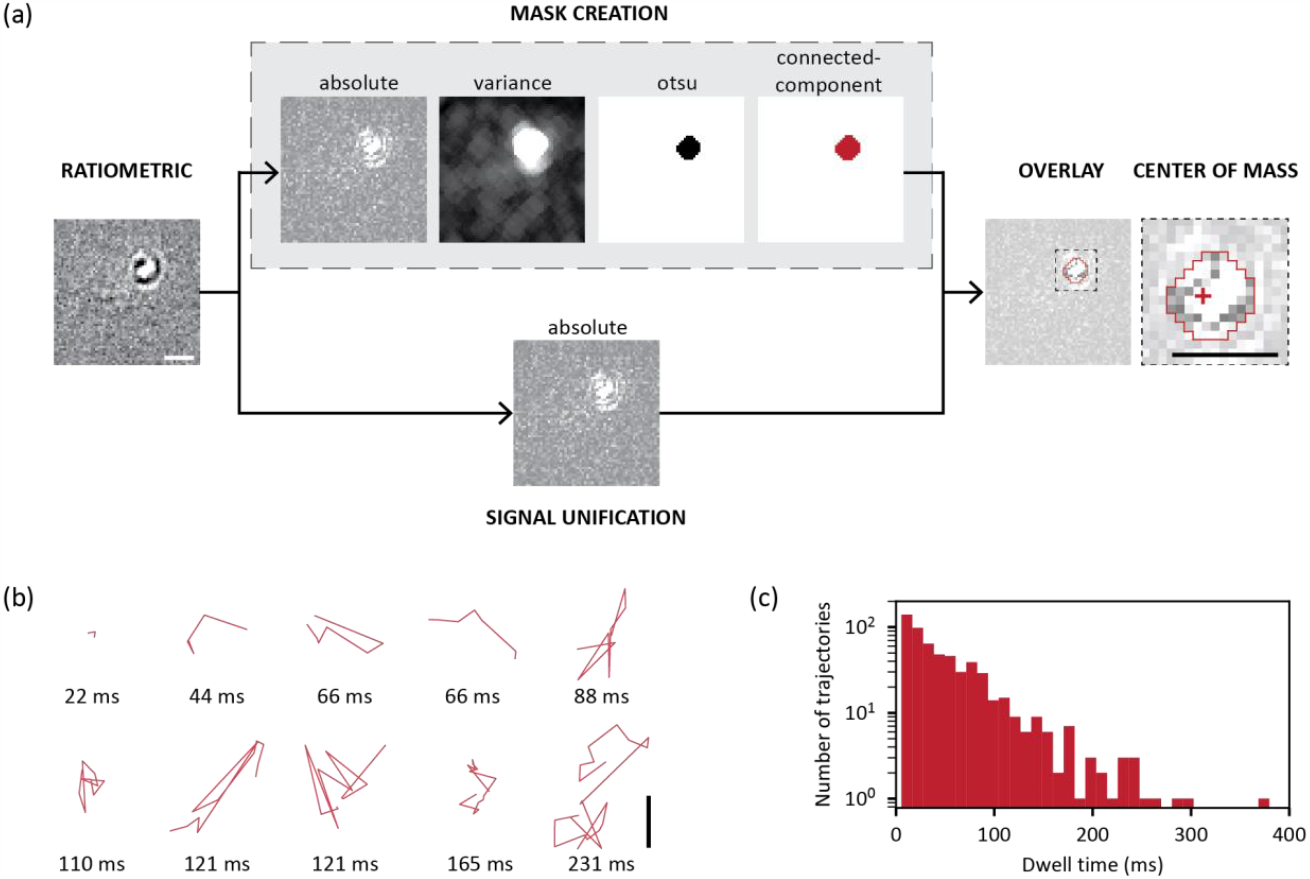
Overview of image processing workflow and of trajectories on pristine hBN. (a) Schematic of image processing workflow. (b) Examples of short and long trajectories on pristine hBN. (c) Dwell time of 20 kbp DNA on pristine hBN. The histogram represents all reconstructed trajectories from three pooled measurement with each 10,000 frames imaged at 90.5 Hz. N (total trajectories) = 581, N (L ≤ 110 ms) = 523, N (L > 110 ms) = 58. Scale bars are 1 µm.

On pristine hBN, we observe a wide range of trajectory geometries and lengths (Figure 2b). We split them into two groups (Figure 2c): those that are shorter than or equal to 110 ms (523 out of 581 total trajectories) and those that are longer than 110 ms (58 out of 581 total trajectories). Overall, the measured mean dwell time for 20 kbp dsDNA on pristine hBN is 51.2 [49.0, 57.4] ms (N= 581 trajectories, *Mean [95% confidence interval*). We noticed some particles diffusing out of the measured FOV (3.4 μm x 3.4 μm), which would lead to a slight underestimate of dwell times. The observed dynamic nature of the DNA trajectories suggests that interactions of dsDNA with the hBN surface are transient, allowing molecules to adsorb temporarily on the surface, and diffuse laterally before detaching and diffusing back into the bulk solution.

Additionally, we noticed a large proportion of brief interactions; in our dataset we measured 554 single-frame binding events, and 581 trajectories that last at least for two frames. Single-frame binding events likely correspond to DNA interacting transiently with the surface due to weak charge-mediated surface interactions. To test this idea, we varied the salt concentration by adding 50 mM of KCl. Under this condition, there appears to be fewer binding events per area and time (Movie S3). Transient binding events, where the DNA molecule diffuses in and out of the imaging volume vs. along the surface, are expected to be short-lived, lasting less than a few milliseconds corresponding to a fraction of a frame. Detecting these events with fluorescence microscopy would be challenging due to their brief duration; however, with iSCAT microscopy, we were able to take advantage of its temporal resolution to capture these events.

We hypothesized that it would be possible to influence the DNA binding kinetics by engineering the surface with fabricated structures that host more defects and hence present more binding sites. To test this idea, we turned to a recently developed surface engineering method^44^ (Figure 3a, SI Experimental Section). While the technique was first demonstrated to produce optical emitters, we saw two potential advantages for our application: there may be minimal damage to the underlying surface of hBN revealed after the water etching step is completed, and the controlled etching may reduce background scattering of the patterned hBN surfaces and edges. As a demonstration, we patterned an array of craters, and monitored the surface with AFM imaging (Figure 3b). The sample was submerged in H_2_O to etch away the patterned surface, and within 15 min craters started forming. We continued imaging the sample until the depth of the craters stabilised at ∼12 nm after about one hour. We also etched lanes using the same process, 1 μm wide, 35 nm deep, and spaced 1 μm apart (Figure 3c). To characterize our structures, we quantified the RMS roughness of the surface. The RMS roughness along the inner trench area of the lanes is 2 nm, while for a single line scan it is even lower (0.3-0.5 nm), and of the surrounding hBN flake area is 1.8 nm (Figure S7).

**Figure 3.**
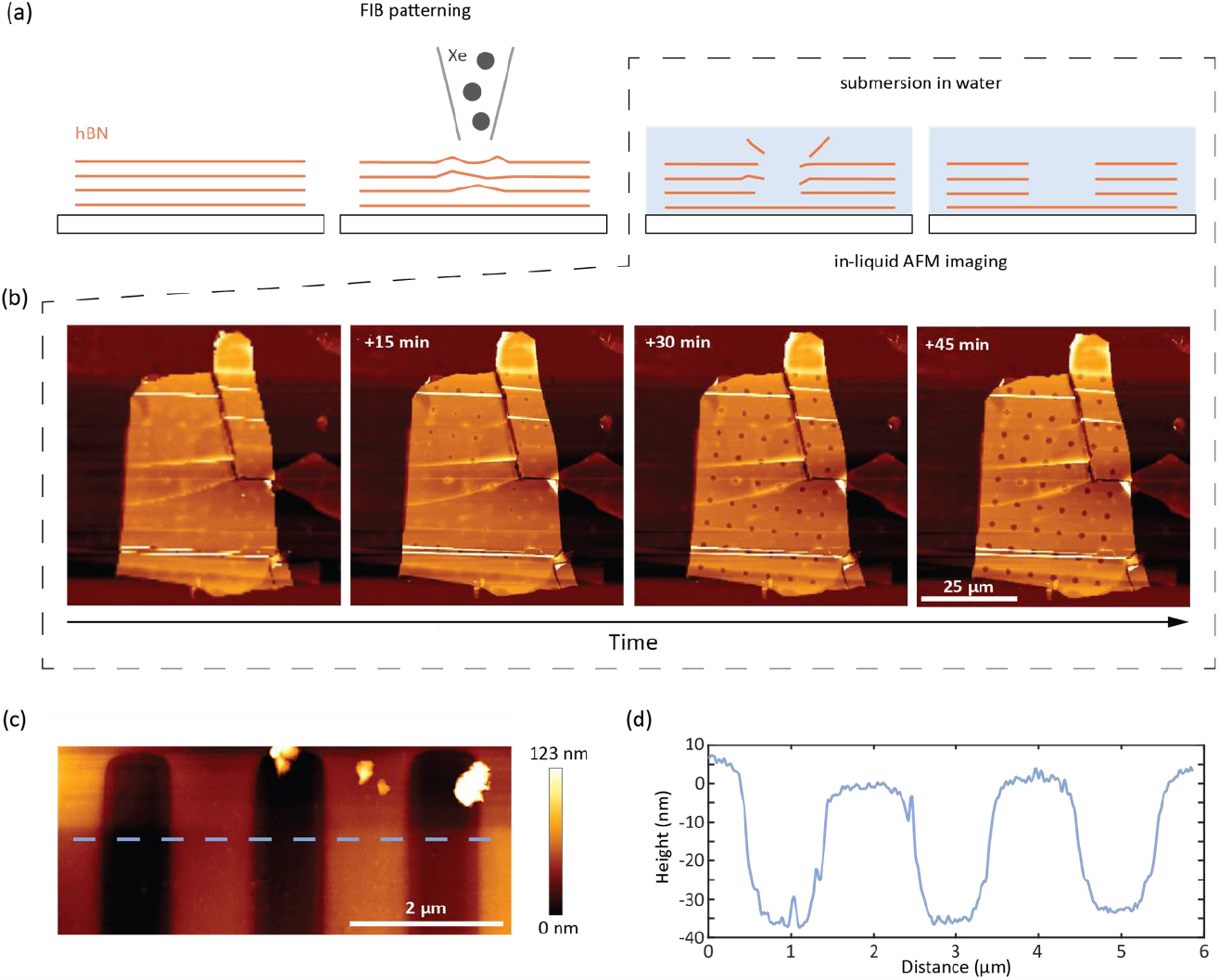
AFM image of pFIB-milled, water-assisted etching of hBN. (a) Illustration of the formation of hBN wells in H_2_O. (b) Time series of a dot-patterned hBN flake upon immersion in water. (c) AFM surface map of tracks milled in hBN flake. (d) AFM height profile scan as indicated by the dashed line in (c), RMS roughness along the trench of the lanes is 2 nm.

To observe the effect of surface modifications on the DNA binding affinity, we performed iSCAT imaging experiments on a single large square FIB-milled onto a flake. Similar FIB milling processes have yielded increased densities of optically active emitters (up to 40x increase, Figure S8) in hBN indicating an increase in defect sites. We reconstructed the trajectories of the DNA on the FIB-milled areas (Figure 4a), and found a mean dwell time of 77.9 [74.8, 86.4] ms (N= 874 trajectories, *Mean [95% confidence interval]*; Figure 4b). This is significantly larger than the dwell time of 51.2 [49.0, 57.4] ms on the native flake. This indicates that there is an increased binding affinity, possibly from interactions with the FIB milling induced defects on the hBN flake. The nanoscopic origins and strength of these interactions are still up for debate but a few hypotheses include increased electrostatic interactions^36^, hydrogen bonding^61^, stacking of the DNA on the hBN^61,62^ and reactivity with chemical species along the defect sites^63^.

**Figure 4.**
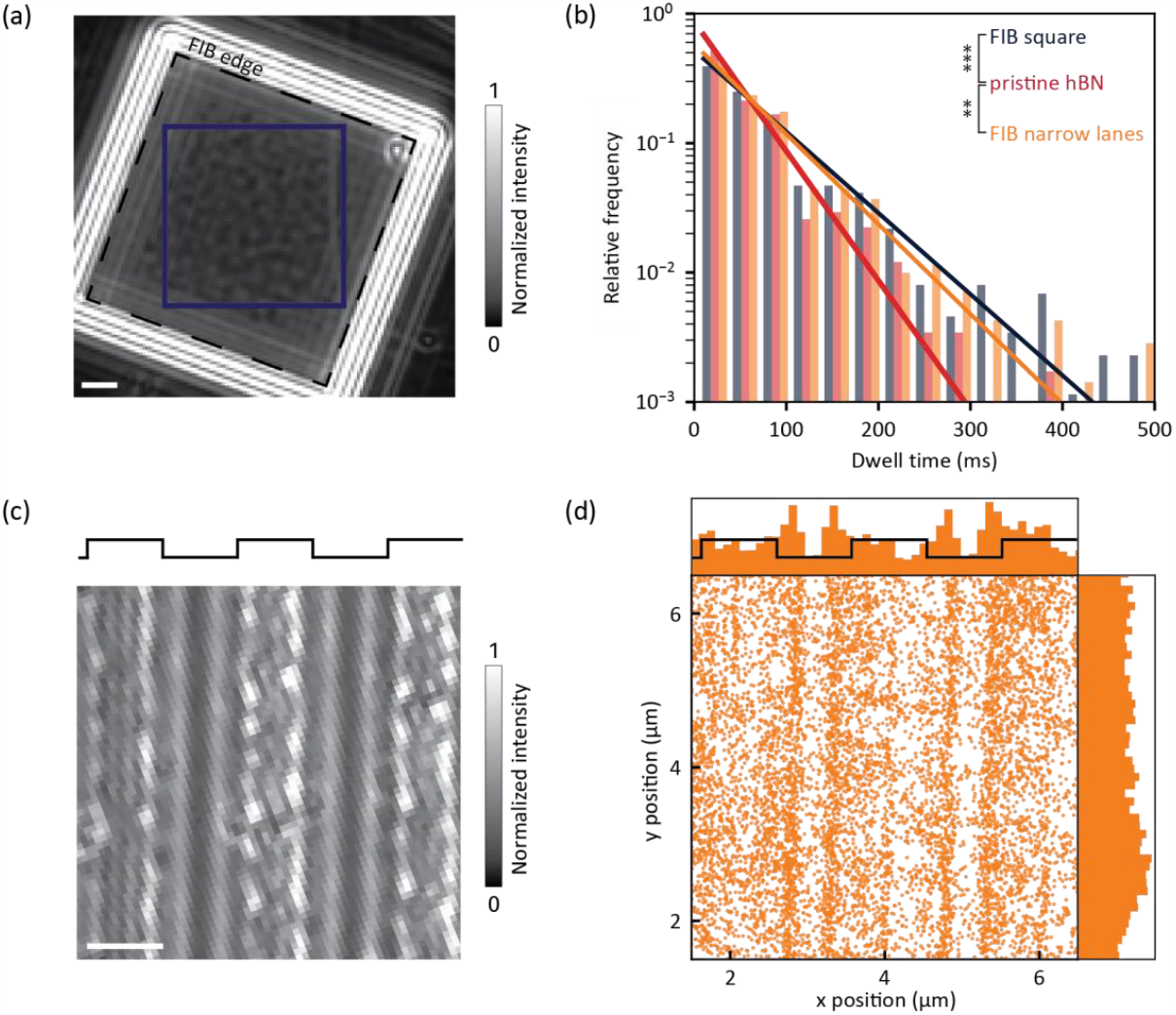
DNA interaction with FIB-milled hBN samples. (a) Native iSCAT image of an hBN flake with an etched square (dashed line). Area used for quantitative analysis is highlighted in blue. (b) Comparison of dwell time of 20 kbp dsDNA on three different surfaces: FIB-etched hBN (FIB square), pristine hBN and hBN patterned with lanes. The histogram represents all reconstructed trajectories from each three pooled measurements. The data was fit to an exponentially decaying distribution. N (FIB square) = 874 trajectories, N (pristine hBN) = 581 and N (FIB lanes) = 703. \*\*\**P*-value = 0.000021, \*\**P*-value = 0.0019. (c) Native iSCAT image of an hBN flake with etched lanes. (d) Distribution of binding spots of 20 kbp dsDNA on sample shown in (c). The scatter histogram represents all localizations from three pooled measurements on the FIB lanes sample shown in (c). N = 9130 localizations. Scale bars are 1 μm.

Next, we performed iSCAT imaging on hBN patterned with narrow lanes by the same process described above for the square (Figure 4c, Movie S2). The edges of the lanes offered scattering contrast, allowing us to contextualize the binding of 20 kbp DNA to the surface. We observed an accumulation of DNA localizations along the edges of the etched lanes (x axis, Figure 4d, Figure S9). The peaks of the accumulation are located between 200-400 nm away from the edges, within the trenches. In contrast, along the direction of the lanes (y axis), binding events appear uniformly distributed. We examined whether scattering from the lane edges could lead to an increased contrast signal for localizations at the edges, increasing the probability of detecting events in those regions and therefore give a false impression of enhanced binding. Further analysis of the mean absolute contrast intensity revealed no such differences in contrast of binding events, and ruled out this possibility (Figure S10). We therefore interpret the data to indicate enhanced binding of DNA to the inside of the lanes. We estimate a DNA polymer radius of gyration of approximately 330 nm, consistent with the location of the peaks displaced from the edges of the lanes.

We wondered whether the preferential binding to lane edges would lead to anisotropic trajectories, e.g. DNA particles preferentially moving along the lanes. To investigate this, we created polar plots of trajectories (Figure S11), and decomposed mean squared displacement (MSD) and step size distributions – along and perpendicular to the lanes (Figure S12, S13, S14). The polar plots do not show a preferential movement along one axis, including when considering only long trajectories. Similarly, the MSD and step size distributions appeared isotropic.

In summary, we present a versatile method based on iSCAT for determining dynamic behavior of DNA on two-dimensional materials, in solution and without labels. We demonstrate that iSCAT can distinguish the binding properties of different interfaces. Additionally, the patterning of 2D material surfaces shows that it is possible to engineer the interaction with dsDNA to create sites with increased interactions. Surprisingly, we did not observe the directed motion of DNA along tracks. This calls for future exploration. Nevertheless, biosensing devices can use engineered interactions to enrich molecules within specific sensing regions^64^ such as the nanopore’s orifice^65,66^, ultimately leading to improved sensitivity in single-molecule sensing^67^.

The data may contain more information about naked DNA’s internal configurations and dynamics. We envision that better control over the background signal and new future analyses incorporating iSCAT PSF modeling or deep-learning^41,68^ will give more insights into these aspects. Since we avoid modification to DNA’s biophysical properties arising from labeling DNA with dye^48^ our method will help to guide the computational modelling of DNA-2D material interactions^36,66^. Our measurements were performed with a temporal resolution of ∼11 ms, but other iSCAT instruments demonstrated a temporal resolution of up 1 MHz^69^. This could help understanding conformational changes in the range of microseconds.

## Acknowledgments

W.Y. and A.R. would like to acknowledge funding from SNF grant CRSII5_193740. This work was supported by the National Centre for Competence in Research (NCCR) Chemical Biology (S.M., J.S.). We thank Lucie Navratilova for help and useful discussions on the plasma FIB milling. We acknowledge Michal Macha for the help with pFIB. We thank Kyle Douglass, the EPFL Hub for Image Analysis and Daniel Sage for their help and useful discussions on the image analysis workflow. We thank Nicolas Hundt for access to and support with the script for median-filtered iSCAT image processing.

## Supporting information

Experimental section and figures as described in the text (PDF)

Supplementary Movie S1: Example of 20 kbp dsDNA on pristine hBN imaged with iSCAT (AVI)

Supplementary Movie S2: Example of 20 kbp dsDNA on FIB-milled hBN imaged with iSCAT (AVI)

Supplementary Movie S3: Example of 20 kbp dsDNA on pristine hBN in DI water and 50 mM KCl (AVI)

## Author contribution

J.S., W.Y., S.M. and A.R. conceived the project and designed experiments. J.S., with assistance of W.Y., performed iSCAT imaging experiments. J.S. implemented the image processing framework and performed data analysis. W.Y. prepared the samples, carried out FIB milling, performed fluorescence imaging with the assistance of Y.S. and N.R. and carried out AFM measurements with assistance from E.M.. J.S., W.Y., S.M. and A.R. designed figures and wrote the manuscript.

