## Supplementary material for "Label-free imaging of DNA interactions with 2D materials": SI

#### Experimental Section

**Reagents, Materials**

The 10, 20 kbp and Lambda DNA fragments were purchased from Thermo Scientific (NoLimits, SM1751 and SM1541, SD0011).

**Sample preparation**

To prepare the samples with focused ion beam (FIB) etching, hexagonal boron nitride (hBN) flakes sourced from high-quality crystals[1] were exfoliated onto glass coverslips (no. 1.5 micro cover glass, Electron Microscopy Sciences, 25 mm in diameter) that had been pre-patterned with gold markers for improved navigation and an electrode mesh to prevent charge buildup. The exfoliation process involved either using tape or gel-pack stamps.

To prepare samples for without FIB etching for interferometric scattering (iSCAT) microscopy measurements, hBN flakes were exfoliated onto a glass coverslip (24 x 50 mm, 0101222, Marienfeld). The cover glass was previously cleaned consecutively for 5 min in the ultrasound bath in each 2% Hellmanex, DI water, isopropanol, DI water, respectively. Between each sonication step three washing steps with DI water were performed.

**iSCAT imaging**

A silicone gasket (3 mm × 1 mm, GBL103250, Grace Bio-Labs) was placed around the region of interest on top of the coverslip to hold the liquid in place.

iSCAT imaging was performed on a Refeyn OneMP setup with a 10.8 µm x 10.8 µm FOV. The laser power was controlled with the unit's internal AOD. The AOD voltage was set to 1V. The samples were placed on the setup and the gaskets filled with 18 µl of Milli-Q water. The focus was locked manually at the edges of the flakes. To image DNA, we added 2 µl DNA and mixed well with a micropipette to achieve a final concentration of 20 ng/ml. The data acquisition typically started within 30 s from adding the DNA. Image stacks were acquired in sets of 10,000 frames with a frame rate of 361.8 Hz. After frame binning, the effective frame rate is 90.5 Hz, resulting in an effective frame length of 11 ms. For subsequent imaging within the same gasket, the focus was set manually again. Multiple sets were acquired for each experimental condition.

**FIB patterning**

The Helios G4 PFIB UXe system equipped with a Xenon Plasma FIB column was utilized for both radiation and hBN patterning. All experiments were conducted at 30 kV, with a 100 pA Xe beam, while adjusting parameters such as dwell time and pitch distance between irradiated spots. The dwell time was set to 500 µs. The typical ion fluence/dose ranged from $1.2 \times 10^{14}$ to $2.5 \times 10^{15}$ ions/cm$^2$, depending on the dwell time.

For this study we used two different patterns: (1) FIB square (a squared area), and (2) FIB narrow lanes (1 µm wide lanes, separated by 1 µm).

**AFM imaging**

To conduct AFM imaging, a customized setup was utilized, comprising a The NanoWizard® 4 XPPK (JPK) positioned above an optical microscope (Olympus IX81). The optical microscope facilitated the detection of the hBN flakes' location in relation to the cantilever. In both air and liquid, AFM images were captured at a line rate of 29 µm/s using TESPA-V2 cantilevers (Antimony doped Si) with a nominal spring constant of 42 N m$^{-1}$ in tapping mode. The cantilever's drive frequency and amplitude were automatically determined through cantilever tuning. The resulting images were processed using standard scanning probe software from JPK.

**Image analysis pipeline for iSCAT data**

The data was analyzed in a custom routine implemted in a Python and ImageJ environment. First, the raw data was drift-corrected with spam[2]. To detect moving DNA molecules, we used an image processing strategy to remove the dominant static background introduced by Heermann et al[3]. Briefly, for the temporally filtered image i, a pixel-wise temporal median image was created from the range of image from i-250 to i+250, so in total 501 images. This temporal median image was divided by image i and subsequently subtracted by 1. This is a continuously running process.

Subsequently, for each temporally filtered image, a binary mask for high-contrast signals was created. This was done in ImageJ[4]. Briefly, a ROI of the image was selected, excluding highly scattering signal from the flake. Then the temporally filtered image was converted to the absolute image, then a variance filter with a radius of 4 px was applied and subsequently an Otsu filter for each individual image was applied.

The binary masks together with the absolute images were imported and overlaid in a custom python script. On the resulting images a connected component analysis was performed and the weighted center of mass (CoM) was calculated based on the pixel values from the absolute image.

Certain restrictions were put in place: (1) Temporally filtered images without a high-contrast signal (presumably without a particle signal) show a high number of white pixels after Otsu filtering. In the case of a 100x100 (60x60) pixel image, a threshold of 400 (150) white pixels per image was introduced. Above this threshold all pixels of the image were set to 0 (black). This was put in place to reduce the number of false-positives. (2) CoMs close to the border of the image (up to 10 pixels = 844 nm) were excluded to prevent border effects.

Trajectories were found with trackpy given the criteria of 15 pixels (= 1.3 µm) radius und 4 frame memory gap (except 0 frame memory gap for MSD and step size distribution calculations).

The detection was validated with by-eye comparison for two conditions: 20 kbp DNA on pristine hBN and 20 kbp DNA on a FIB lanes sample. In the case of 20 kbp DNA on pristine hBN, we compared 200 frames by eye. By eye, 60 localizations were observed, whereas the automated pipeline detected 48. In total, 45 detections were true-positive, leading to a detection rate of 75 %. In the case of 20 kbp DNA on pristine hBN, we compared 50 frames by eye. By eye, 54 localizations were observed, whereas the automated pipeline detected 42. In total, 33 detections were true-positive, leading to a detection rate of 61 %.

**Trajectory duration estimations**

For trajectory duration estimations, the data from each three measurements per sample/ condition was pooled and was fitted to an exponentially decaying distribution in Python. For each condition, the mean fitted value and the 95 % confidence interval was reported. To ensure the identification of dynamic DNA molecules, we implemented the following definition: A particle requires to be detected in two (consecutive) frames to be assigned a dwell time of a frame length (11 ms; so we assume an actual dwell time of the particle from 5.5 to 16.5 ms).

Statistical significance between datasets was determined by Kolmogorov- Smirnov tests in Python (SciPy). *P*-values of $< 0.05$ were used to reject the null hypothesis.

#### Figure S1. Scattering of hBN flake edges in iSCAT images

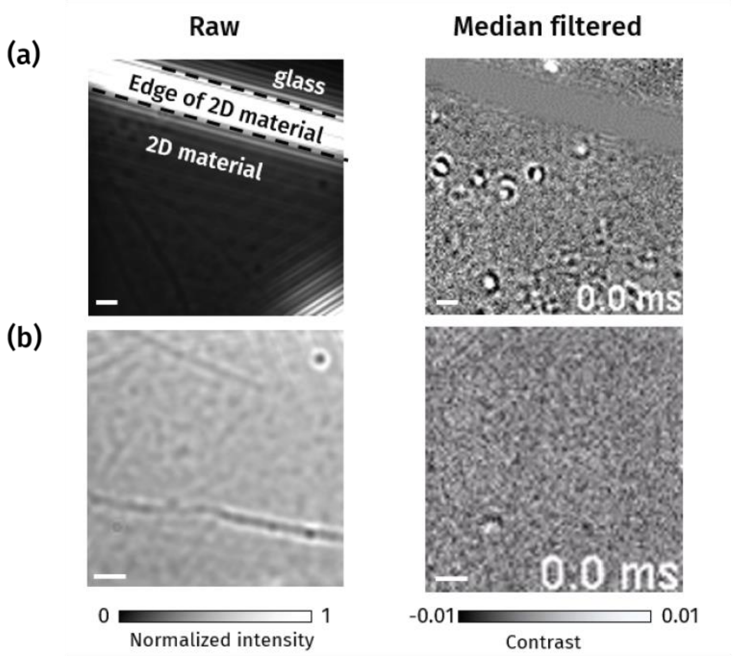

**Figure S1.** Scattering of hBN flake edges in iSCAT images. (a) Example of an hBN flake imaged with its edge in the FOV. (b) Example of an hBN flake imaged in the center of the flake, without any edges in the FOV. Scale bars are 1 µm.

#### Figure S2. Time series of median-filtered images of DNA molecules of different lengths

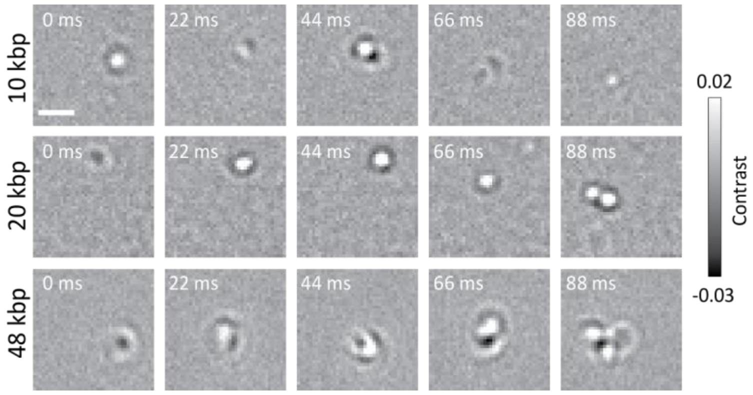

**Figure S2.** Time series of median-filtered images of DNA molecules of different lengths.

#### Figure S3. Time series of mean-filtered images

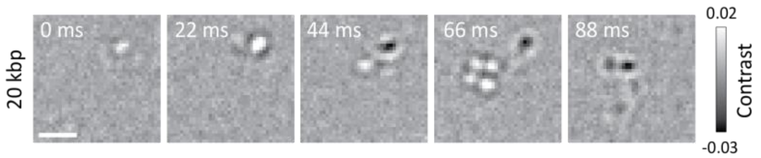

**Figure S3.** Time series of mean-filtered images. The images show the same 20 kbp DNA particle and ROI as shown in Figure S2. Scale bar is 1 µm.

### Figure S4. Parameters used for quantitative analysis of iSCAT data

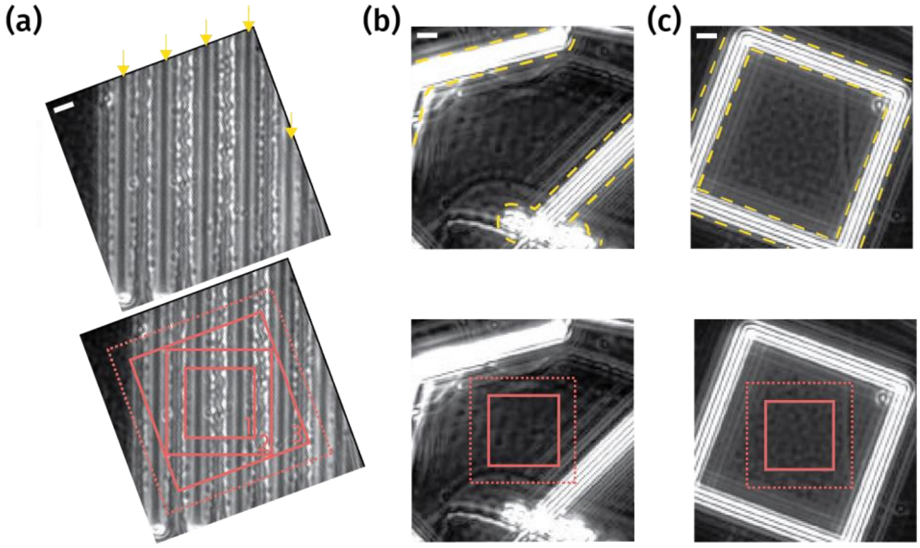

| Date | M ID | Pattern | Temporal filtering | Drift correction | Size of ROI used for image pre-processing [pixels$^2$] | Size of ROI used for CoM analysis [pixels$^2$] | Tracking parameters | Used for figure |
|---|---|---|---|---|---|---|---|---|
| 221018 | 30 | FIB lanes | Median, 501 | Spam | 100x100, centered | 80x80 | radius 15 pixels, memory gap 4 frames | Fig S9a, Fig S10, Fig S11a, Fig S12 |
| 221018 | 31 | FIB lanes | Median, 501 | Spam | 100x100, centered | 80x80 | radius 15 pixels, memory gap 4 frames | Fig S9a, Fig S10, Fig S11a, Fig S12 |
| 221018 | 32 | FIB lanes | Median, 501 | Spam | 100x100, centered | 80x80 | radius 15 pixels, memory gap 4 frames | Fig S9a, Fig S10, Fig S11a, Fig S12 |
| 221018 | 30 | FIB lanes | Median, 501 | Spam | 100x100, centered | 80x80 | radius 15 pixels, memory gap 0 frames | Fig S13a, Fig S14a |
| 221018 | 31 | FIB lanes | Median, 501 | Spam | 100x100, centered | 80x80 | radius 15 pixels, memory gap 0 frames | Fig S13a, Fig S14a |
| 221018 | 32 | FIB lanes | Median, 501 | Spam | 100x100, centered | 80x80 | radius 15 pixels, memory gap 0 frames | Fig S13a, Fig S14a |
| 221018 | 30 | FIB lanes | Median, 501 | Spam | 100x100, centered | 60x60, rotated by -68 deg | radius 15 pixels, memory gap 4 frames | Fig 4d |
| 221018 | 31 | FIB lanes | Median, 501 | Spam | 100x100, centered | 60x60, rotated by -68 deg | radius 15 pixels, memory gap 4 frames | Fig 4d |
| 221018 | 32 | FIB lanes | Median, 501 | Spam | 100x100, centered | 60x60, rotated by -68 deg | radius 15 pixels, memory gap 4 frames | Fig 4d |
| 221018 | 30 | FIB lanes | Median, 501 | Spam | 100x100, centered | 40x40, rotated by -68 deg | radius 15 pixels, memory gap 4 frames | Fig 4b, Fig S9b, Fig S11b |
| 221018 | 31 | FIB lanes | Median, 501 | Spam | 100x100, centered | 40x40, rotated by -68 deg | radius 15 pixels, memory gap 4 frames | Fig 4b, Fig S9b, Fig S11b |
| 221018 | 32 | FIB lanes | Median, 501 | Spam | 100x100, centered | 40x40, rotated by -68 deg | radius 15 pixels, memory gap 4 frames | Fig 4b, Fig S9b, Fig S11b |
| 221018 | 35 | pristine hBN | Median, 501 | Spam | 60x60, centered | 40x40 | radius 15 pixels, memory gap 4 frames | Fig 2b,c, Fig 4b, Fig S9c, Fig S11c |
| 221018 | 36 | pristine hBN | Median, 501 | Spam | 60x60, centered | 40x40 | radius 15 pixels, memory gap 4 frames | Fig 2b,c, Fig 4b, Fig S9c, Fig S11c |
| 221018 | 37 | pristine hBN | Median, 501 | Spam | 60x60, centered | 40x40 | radius 15 pixels, memory gap 4 frames | Fig 2b,c, Fig 4b, Fig S9c, Fig S11c |
| 221018 | 35 | pristine hBN | Median, 501 | Spam | 60x60, centered | 40x40 | radius 15 pixels, memory gap 0 frames | Fig S13b, Fig S14b |
| 221018 | 36 | pristine hBN | Median, 501 | Spam | 60x60, centered | 40x40 | radius 15 pixels, memory gap 0 frames | Fig S13b, Fig S14b |
| 221018 | 37 | pristine hBN | Median, 501 | Spam | 60x60, centered | 40x40 | radius 15 pixels, memory gap 0 frames | Fig S13b, Fig S14b |
| 221018 | 40 | FIB square | Median, 501 | Spam | 60x60, imagej position (30,37) | 40x40 | radius 15 pixels, memory gap 4 frames | Fig 4b, Fig S9d, Fig S11d |
| 221018 | 41 | FIB square | Median, 501 | Spam | 60x60, imagej position (30,37) | 40x40 | radius 15 pixels, memory gap 4 frames | Fig 4b, Fig S9d, Fig S11d |
| 221018 | 42 | FIB square | Median, 501 | Spam | 60x60, imagej position (30,37) | 40x40 | radius 15 pixels, memory gap 4 frames | Fig 4b, Fig S9d, Fig S11d |

**Figure S4.** Parameters used for quantitative analysis of iSCAT data. The top row shows highly scattering areas, the bottom row the ROIs used for quantitative analysis (dotted lines for analysis up to otsu filter, solid lines for calculation of center of mass (CoM, binding spots)). (a) Etched lanes sample: The etched lanes are highlighted with yellow arrows. In the bottom row, the large dotted line refers to the cropped area used in the image analysis pipeline up to the otsu filter (100 x 100 pixel$^2$). The solid lines refer to different ROIs used for calculating the CoMs. The solid line (3) is used for trajectories and large ROI binding histograms (80 x 80 pixel$^2$, SI), (2) for the binding histogram in the main text (60 x 60 pixel$^2$) and (1) for comparing the same sized ROI with examples in (b) and (c) (40 x 40 pixel$^2$). (b) Pristine hBN sample: The top image shows highly scattering areas from presumably cracks within in the flake. (c) Etched square sample: The top image shows the etched edges of the square pattern. The images shown are drift-corrected (SI Experimental Section). Scale bars are 1 µm. The table shows all parameters used for analysis.

### Figure S5. Detailed overview of image processing workflow

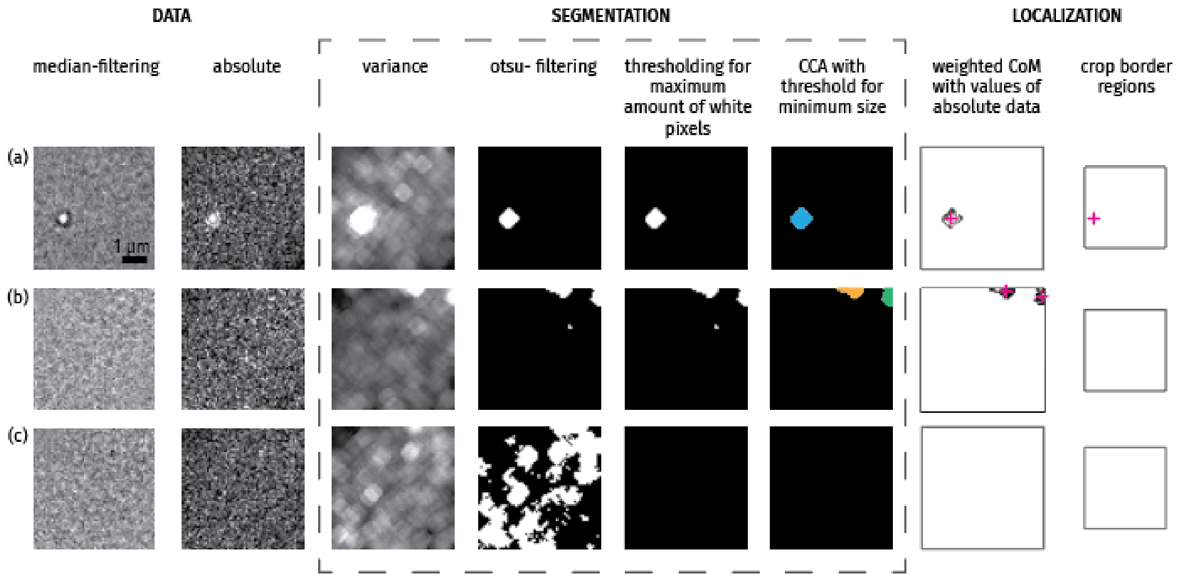

**Figure S5.** Detailed overview of image processing workflow. Example frames taken from movie of 20 kbp dsDNA in DI water. (a) Frame with one high-intensity signal. (b) Frame with small high-intensity signal at the border of the recorded FOV. (c) Frame with no high-intensity signal.

#### Figure S6. Effect of variance filter in image analysis pipeline

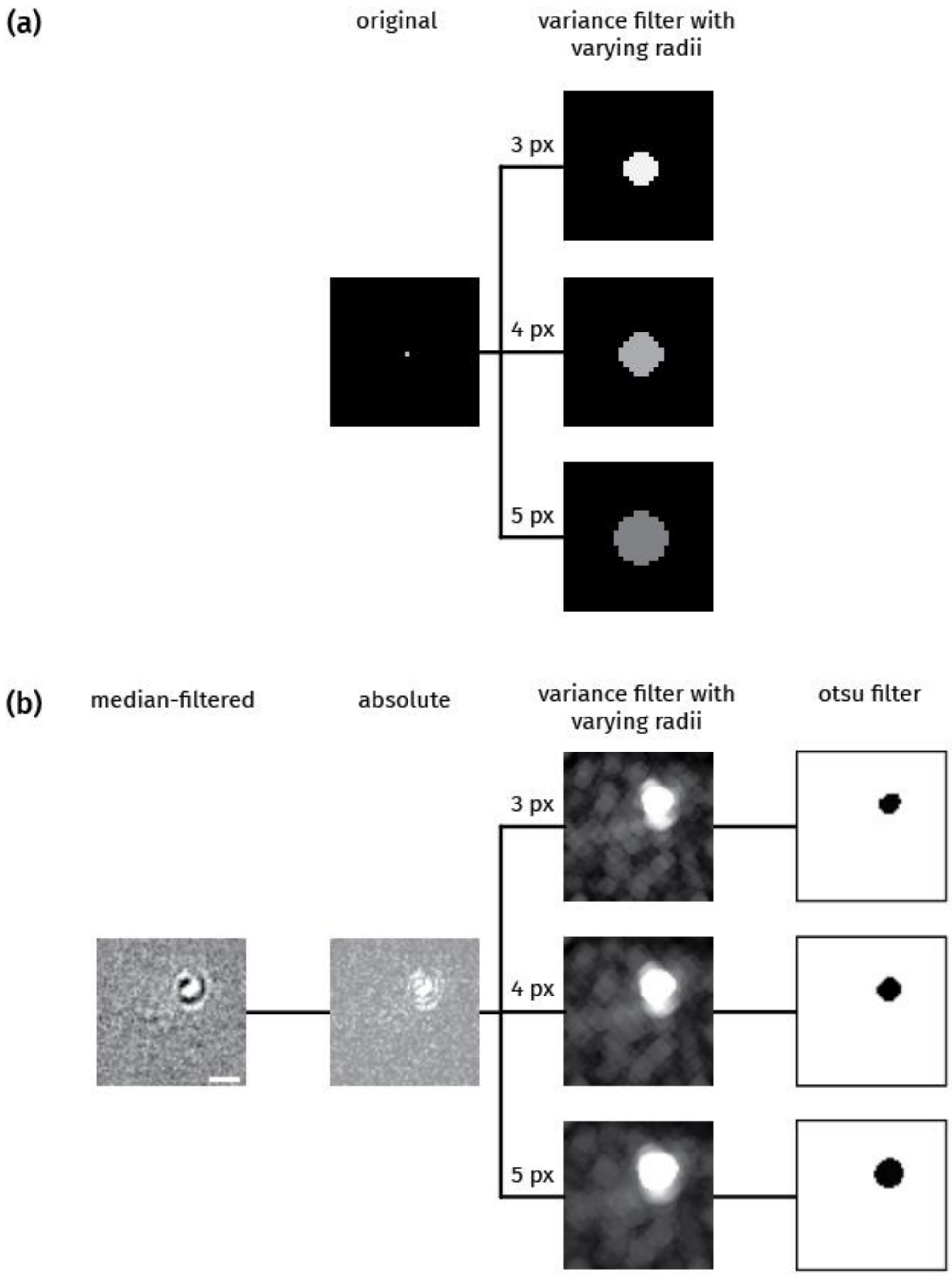

**Figure S6.** Effect of variance filter in image analysis pipeline. (a) Example of the effect of the variance filter with varying radii (3 pixel, 4 pixel, 5 pixel) applied to an artifical 30 x 30 pixel$^2$ image (all pixel values are set to 0, center pixel set to 50). (b) Example of the effect of the variance filter with varying radii (3 pixel, 4 pixel, 5 pixel) on the example image shown in Figure 2a. Scale bar is 1 µm.

#### Figure S7. Atomic force microscopy (AFM) profile of FIB milled lanes

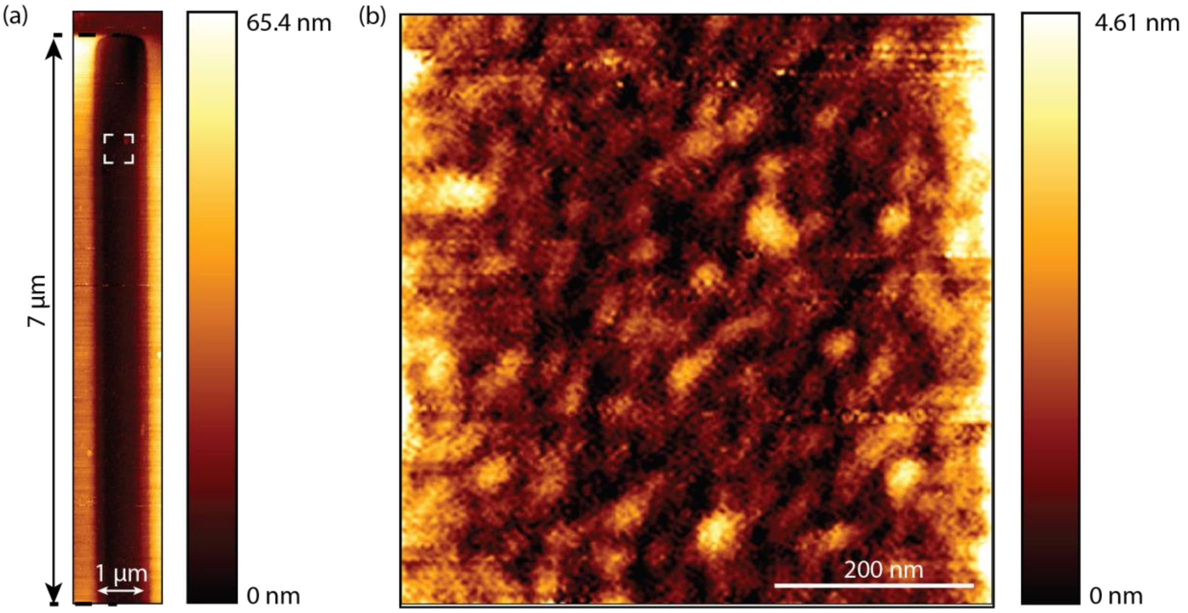

**Figure S7.** Atomic force microscopy (AFM) profile of FIB-milled lanes fabricated using pFIB. (a) AFM image of one of the lanes. (b) High-resolution scan of the inner lane surface (area marked by white square in (a)).

#### Figure S8. Optically active defects in hBN flake induced by pFIB

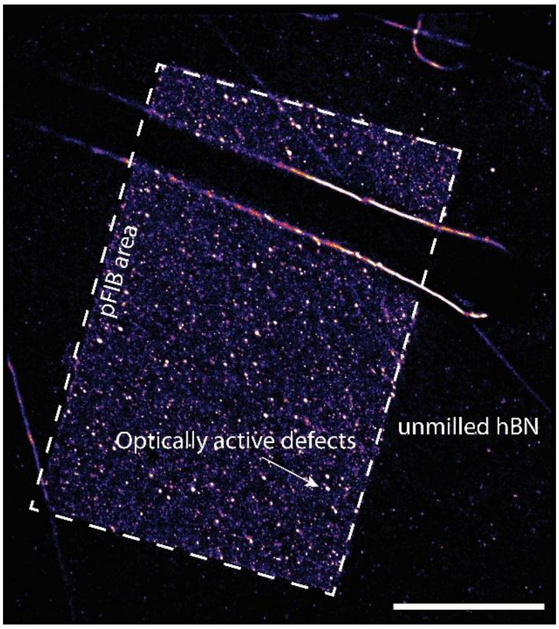

**Figure S8.** Optically active defects in hBN flake induced by pFIB. Optically active defects excited by 561 nm laser and localised by single-molecule localisation microscopy after pFIB. We compare two areas, an area exposed to the pFIB and a control area of the flake next to it. The area that has been irradiated shows an increase in optical activation. We find an average of 613 emitters per µm$^2$ in the FIB area and 14.4 emitters per µm$^2$ in the non-FIB area upon acquisition of 5,000 frames. Scale bar indicates 10 µm.

### Figure S9. Binding spots of 20 kbp DNA on different hBN samples

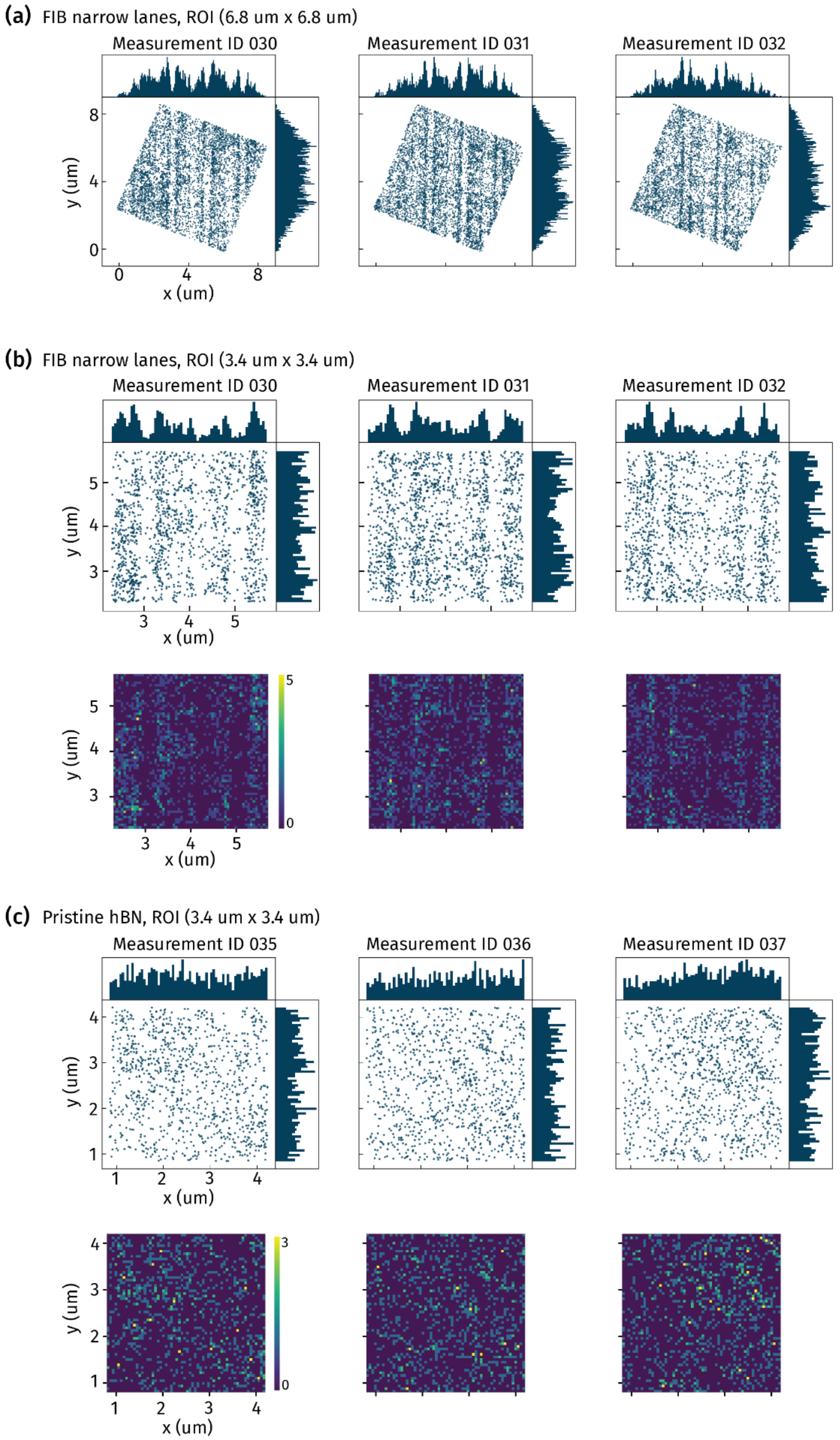

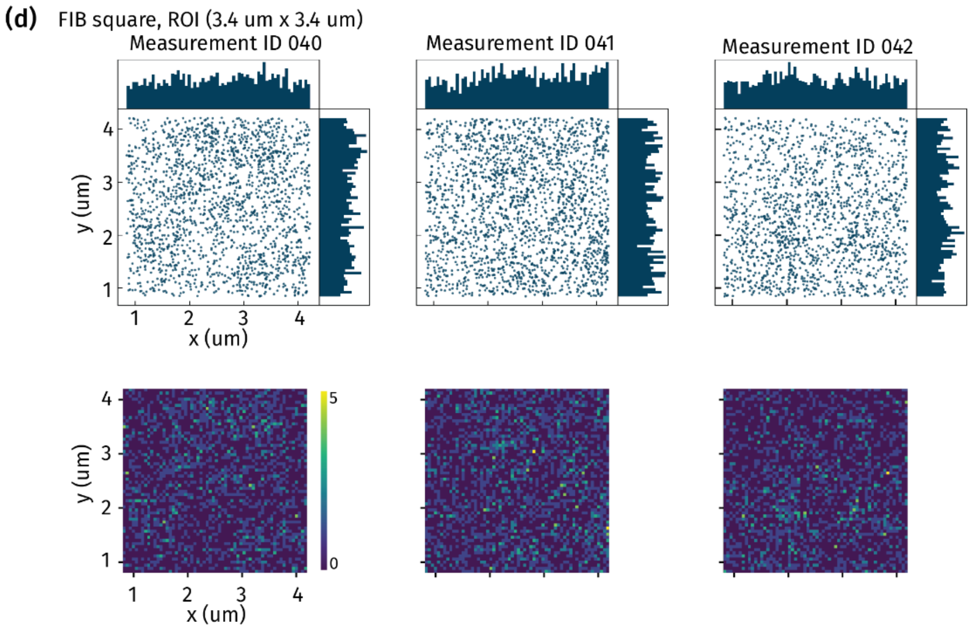

**Figure S9.** Binding spots of 20 kbp DNA on different hBN samples. (a) Binding spots on a large ROI with etched lanes (1 µm wide and 1 µm apart), N (MID 030, 031, 032) = 5746, 6149, 5214. (b) Cropped ROI from (a), cropping limits: 2.3 and 5.7 µm, N (MID 030,031,032) = 1459, 1575, 1374. (c) Binding spots on pristine hBN, N (MID 035, 036, 037) = 991, 969, 1026. (d) Binding spots on completely etched hBN surface (without pattern in ROI), N (MID 040, 041, 042) = 1821, 2055, 1677. The heatmaps were generated with 60 bins per axis (width per bin 0.057 um). All measurements were recorded with a frame rate of 90.5 Hz up to 10,000 frames.

### Figure S10. Comparison of mean absolute contrast intensity of localizations close ('edge') and further away ('outside') to the edges in patterned hBN with 20 kbp dsDNA

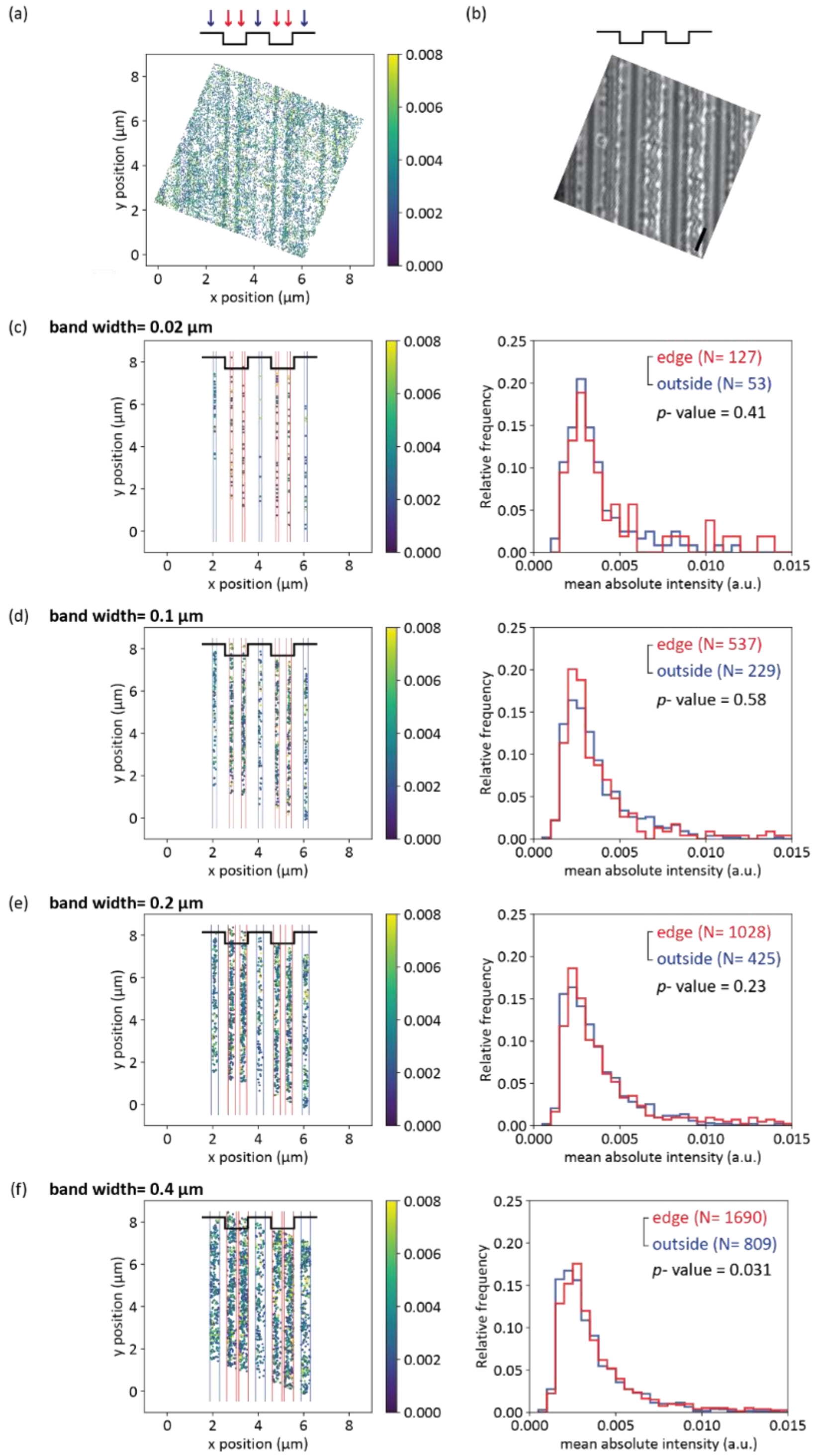

**Figure S10.** Comparison of mean absolute contrast intensity of localizations close ('edge') and further away ('outside') to the edges in patterned hBN with 20 kbp dsDNA. a) Mean absolute intensity of all detected localization of the FOV. (b) Native iSCAT image. (c- f) Localizations considered for the analysis and their respective distribution of mean absolute intensities. P- values are based on a Kolmogorov-Smirnov test.  Scale bar is 1 µm.

### Figure S11. Polar plots of reconstructed trajectories of 20 kbp DNA

**(a)** FIB narrow lanes, ROI (6.8 um x 6.8 um), memory gap 4

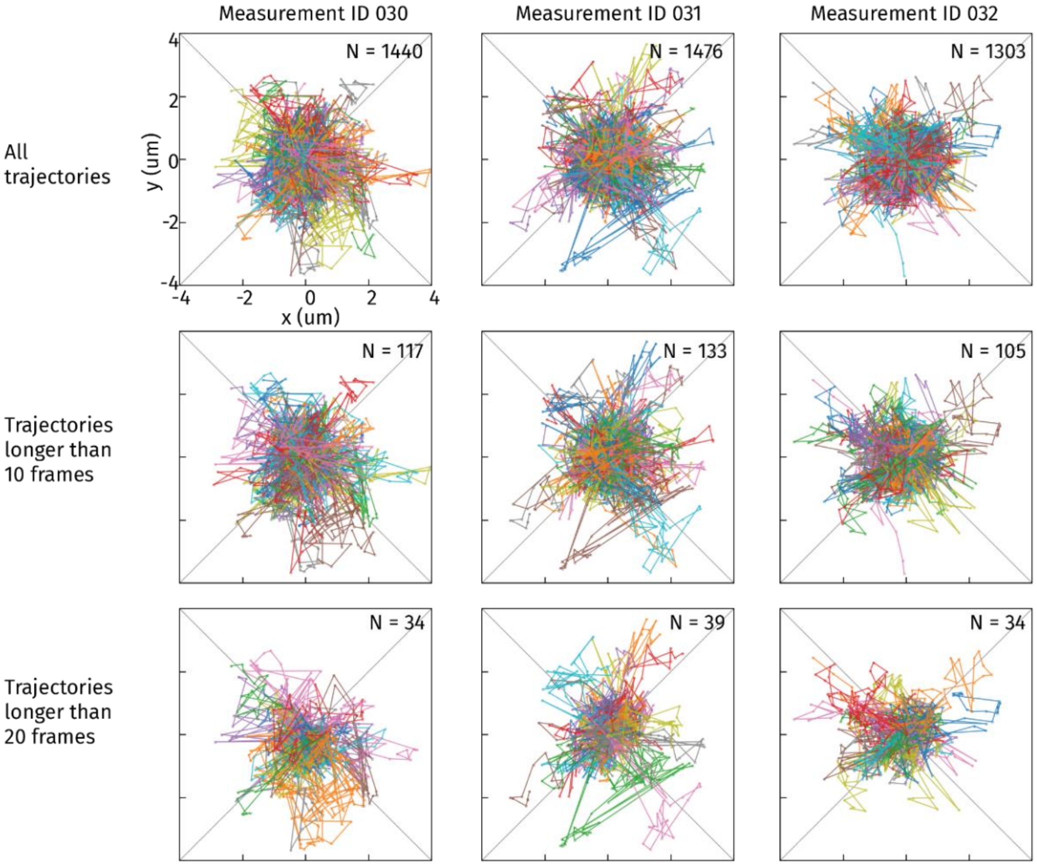

**(b)** FIB narrow lanes, ROI (3.4 um x 3.4 um), memory gap 4

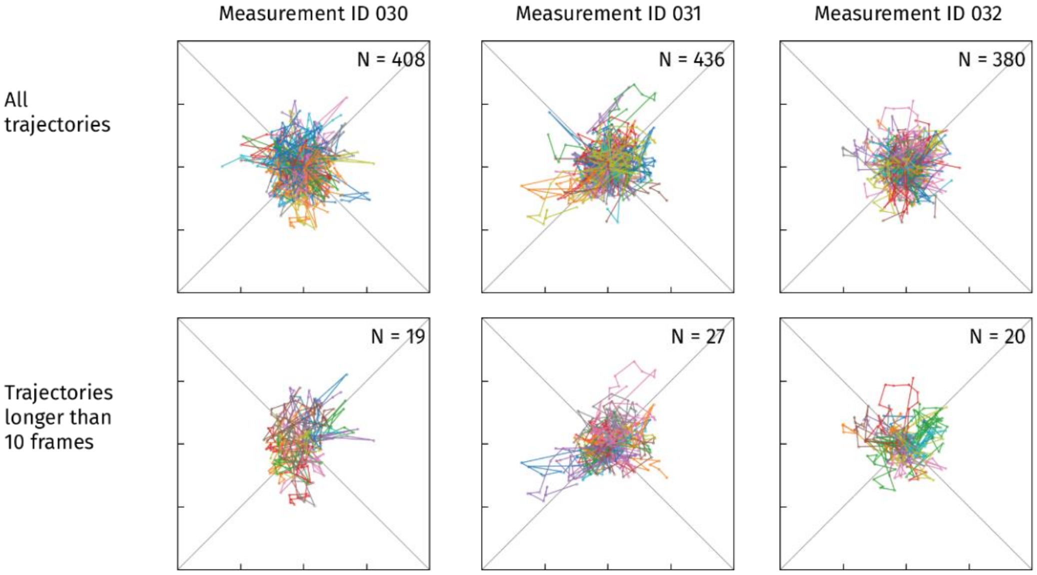

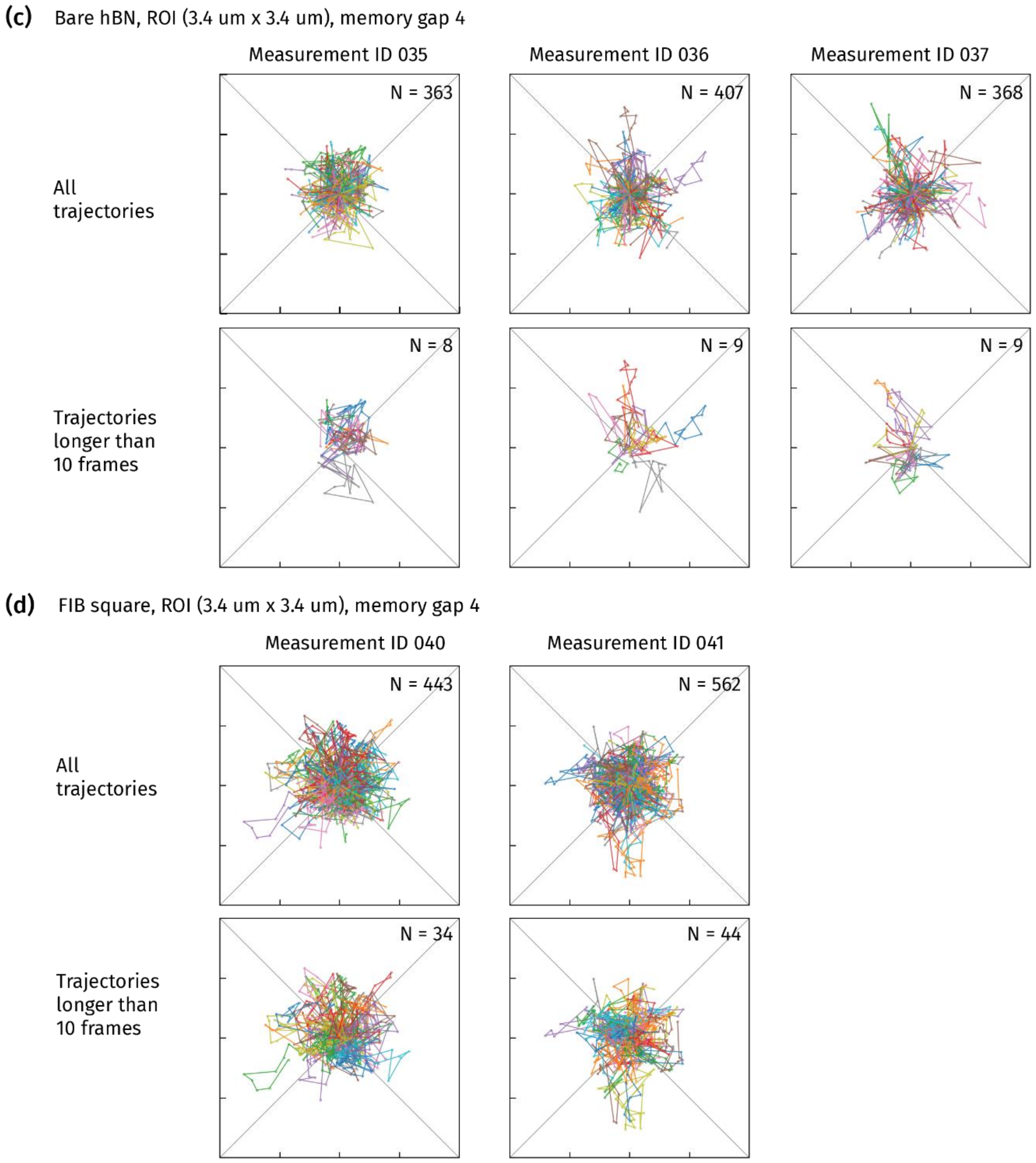

**Figure S11.** Polar plots of reconstructed trajectories of 20 kbp on different FIB structures on hBN. All measurements were recorded with a frame rate of 90.5 Hz up to 10,000 frames.

### Figure S12. Reconstructed trajectories of 20 kbp DNA on FIB-milled hBN samples

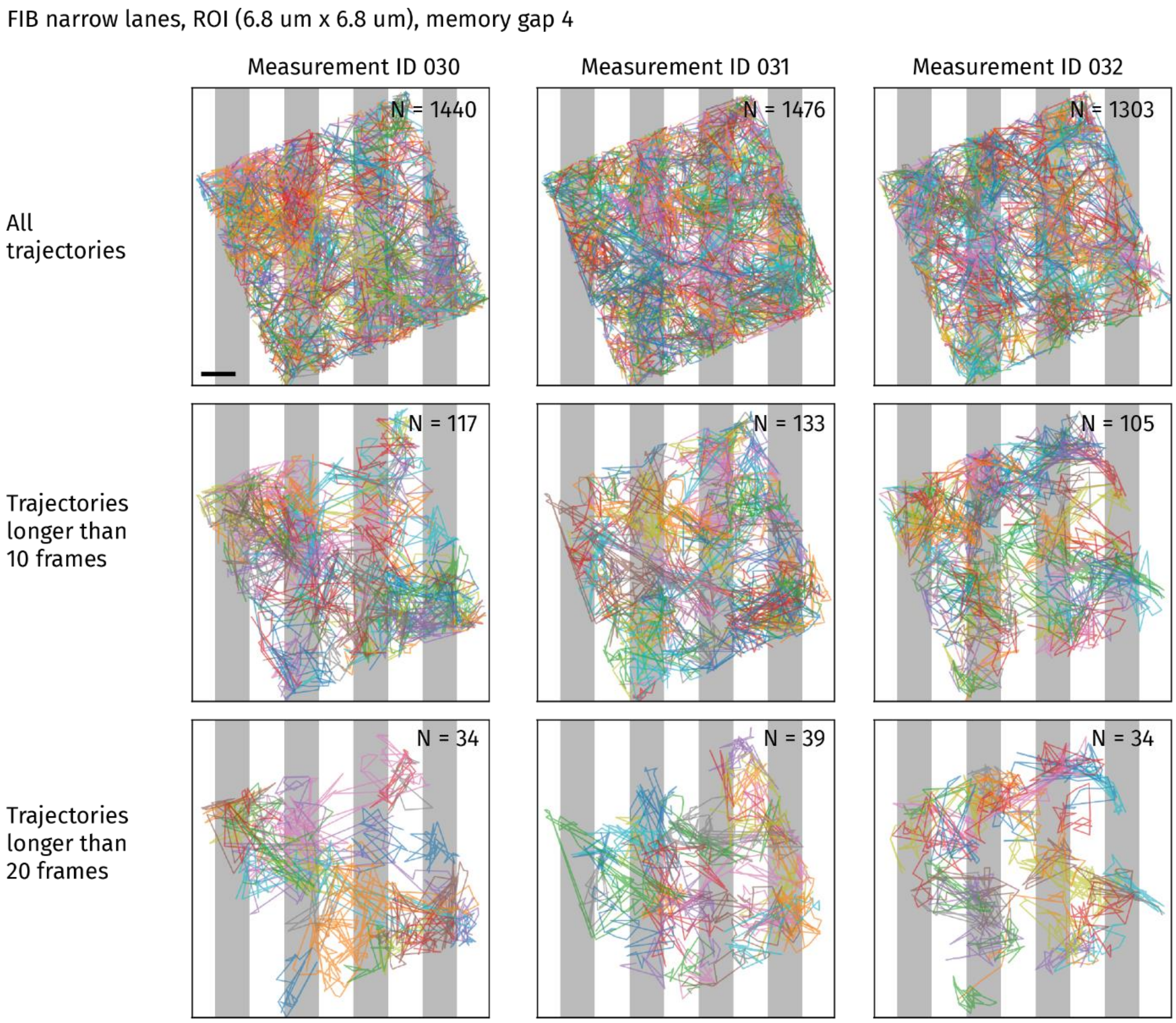

**Figure S12.** Reconstructed trajectories of 20 kbp DNA on FIB-milled hBN samples. The grey areas are schematics and represent the bottom of the lanes. Scale bar is 1 µm.

### Figure S13. Decomposed mean squared displacement (MSD) of trajectories of 20 kbp DNA on hBN imaged with iSCAT

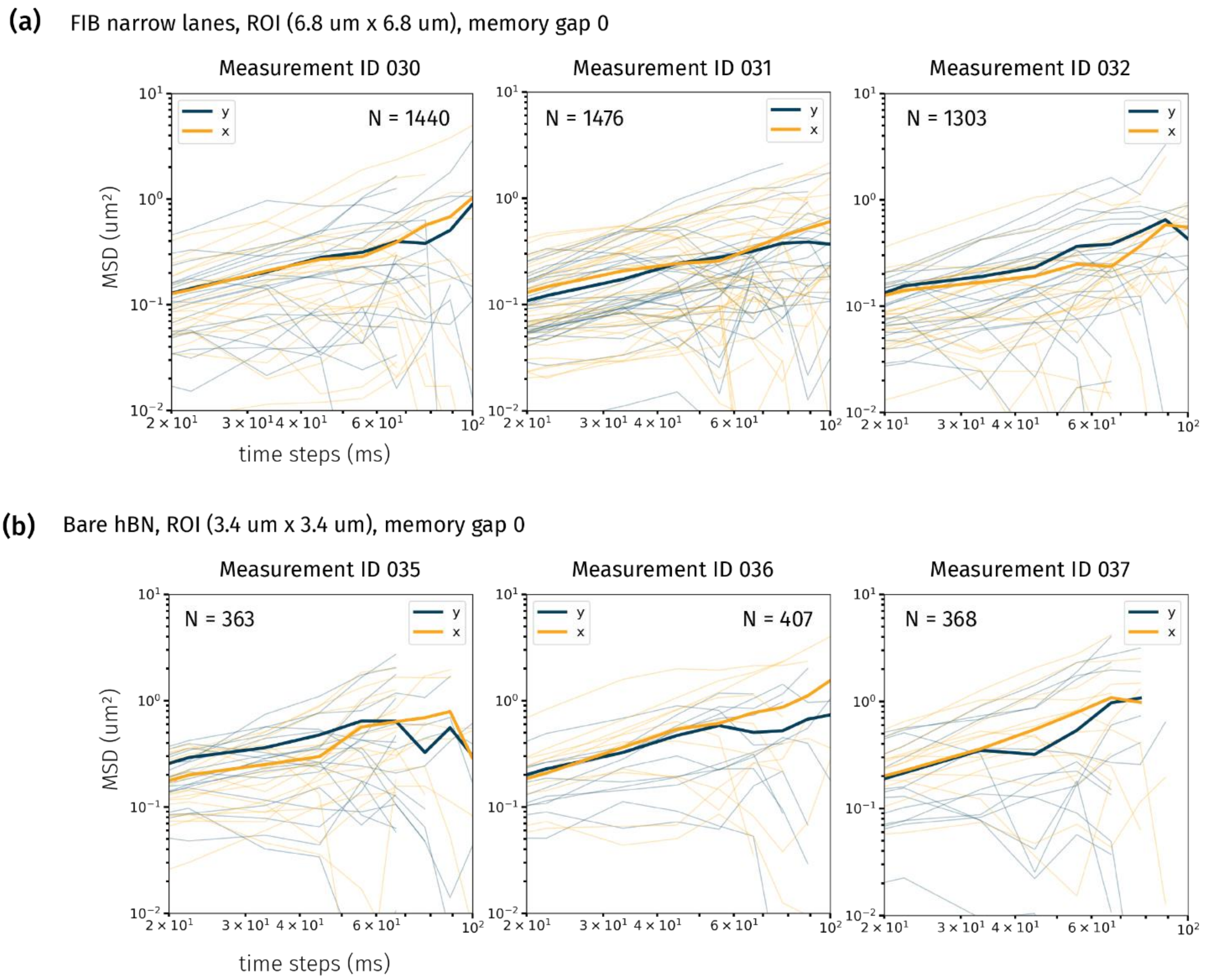

**Figure S13.** Decomposed mean squared displacement (MSD) of trajectories of 20 kbp DNA on hBN imaged with iSCAT. (a) On etched lanes, analyzed with large ROI. The data in (a) was turned by -68 degree, so that the x axis is perpendicular and the y axis parallel to the etched lanes. (b) On pristine hBN. For this analysis the trajectories were reconstructed with a memory gap of 0 frames (SI Experimental Section).

### Figure S14. Decomposed step displacement distribution of trajectories of 20 kbp DNA on hBN imaged with iSCAT

**(a)** FIB narrow lanes, ROI (3.4 um x 3.4 um), memory gap 0

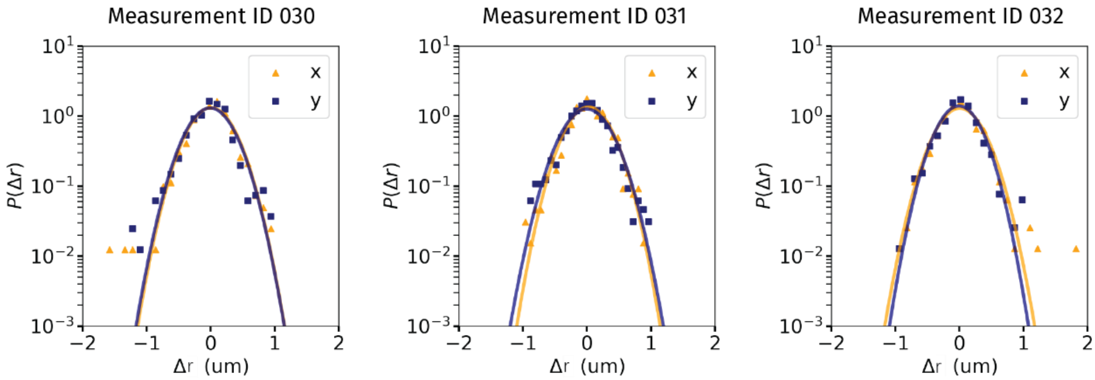

**(b)** Bare hBN, ROI (3.4 um x 3.4 um), memory gap 0

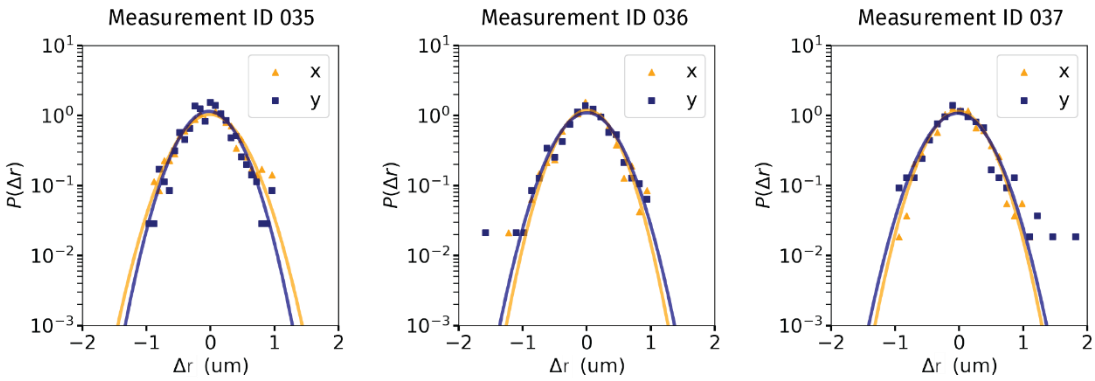

**Figure S14.** Decomposed step displacement distribution of trajectories of 20 kbp DNA on hBN imaged with iSCAT. (a) On etched lanes, analyzed in small ROI. (b) On pristine hBN. The data (represented by triangles and squares) was fitted to a Gaussian distribution (represented by a line). For this analysis the trajectories were reconstructed with a memory gap of 0 frames (SI Experimental Section).
